# Build a better mouse task – can an open-source rodent joystick enhance reaching behavior outcomes through improved monitoring of real-time spatiotemporal kinematics?

**DOI:** 10.1101/560961

**Authors:** Parley Belsey, Mark A. Nicholas, Eric A Yttri

## Abstract

For decades, advanced behavioral tasks have only been used in human and non-human primates. However, with improved analytical and genetic techniques, there has been a growing drive to implement complex reaching, decision-making, and reaction time tasks – not in primates – but in rodents. Here, we assess the hypothesis that a mouse can learn a cued reaction time task. Moreover, we tested multiple training regimens and found that introducing elements of the reaction time task serially hindered, rather than helped task acquisition. Additionally, we include a step-by-step manual for inexpensive implementation and use of a rodent joystick for behavioral analysis. Task and analysis code for the evaluated behaviors are included such that they may be replicated and tested further. With these, we also include code for a probabilistic reward ‘two-arm bandit’ task. These various tasks, and the method to construct and implement them, will enable greatly improved study of the neural correlates of behavior in the powerful mouse model organism. In summary, we have tested and demonstrated that mice can learn sophisticated tasks with A joystick, and that targeted task design provides a significant advantage. These results of this study stand to inform the implementation of other sophisticated tasks using the mouse model.

## INTRODUCTION

The study of behavioral spatiotemporal dynamics in the mouse has grown considerably in recent years (Harvey et al., 2009; Cohen et al., 2015; Guo et al., 2015; Klaus et al., 2017), opening new avenues of discovery that take advantage of the optogenetic advantages the mouse model provides. Rodent work enables high-throughput methods that accelerate discovery and capture behavioral variance, rather than ignore it (Brunton et al., 2013; Guo et al., 2015). The growth of rigorous mouse behavioral paradigms also reduces barriers to entry that may limit smaller labs and institutions. Therefore, it is advantageous for the field to establish inexpensive methods to study highly-quantifiable behavior in an awake behaving mouse (Fetsch, 2016).

Reaching is a well-studied behavior across several species (Fromm and Evarts, 1981; Churchland et al., 2012; Dean et al., 2012; Cherian et al., 2013; Yttri et al., 2013; Mathis et al., 2017). This goal-oriented behavior is a unitary, highly-quantifiable movement, whereas other tasks require several actions, like reorientation followed by locomotion across a cage (Tai et al., 2012; Lak et al., 2014;). Despite this, the behavior provides rich spatiotemporal dynamics (Bollu et al., 2018) that do not exist in other presses. Joysticks have been used for decades with both human and nonhuman primates(Thoroughman and Shadmehr, 1999; Maeda et al., 2018), and more recently with rats(Slutzky et al., 2010), and can provide a real-time readout of the X and Y trajectory. In obtaining position and speed information in real time, joysticks enable the study of ongoing correlated neural activity (Paninski, 2003; Panigrahi et al., 2015) or stimulation in closed loop triggered off a specific spatiotemporal feature of movement (Yttri and Dudman, 2016). This feature presents a significant advantage over impressive, but post-hoc, motion capture techniques (Guo et al., 2015; Mathis et al., 2018; Robie et al., 2017) – though computer vision methods are quickly advancing real-time capabilities (Ellens et al., 2016).

## METHODS

Adult (>p40) wild-type C57/Bl6 mice were used. All animals were water-deprived, but kept at least 75% of their free weight. Before training, animals were accustomed to experimenter handling and head fixation (see Osborne and Dudman, 2014) via a surgically implanted head cap. Three conditions were considered. First, animals were trained to reach in any direction past an amplitude threshold for reward. Reward amplitude threshold was a function of the animal’s performance success. This condition was common across all conditions through to session 13 (Figure 1, red line), and was the only requisite of the task in the CONTROL training regimen.

The reaction time conditions were either LIGHT EARLY or LIGHT LATE. In the LIGHT EARLY Condition, the go cue light was introduced on the first day, although no punishment (a time out and house lights coming on) occurred until day 14. The LIGHT LATE Condition introduced the light in the seventh session. Two animals were used in each condition, with equal numbers of male and female mice used. As our N is quite small in this documentation of our preliminary original research findings, all statistics - including error bars - have been left out. All procedures were approved by the *Institutional Animal Care and Use Committee* at Carnegie Mellon University.

**Figure 1:**
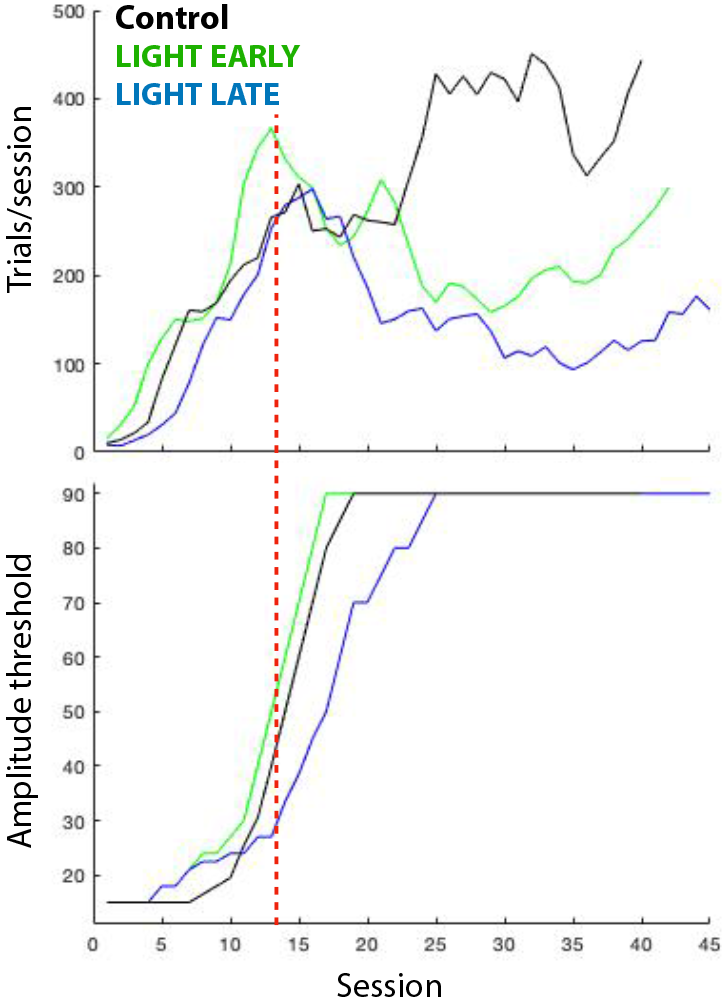
Effects of different training regimens. Average number of trials performed by mice in each session (top) and the amplitude of the reward threshold of those sessions. Reward amplitude threshold, the distance a reach is required to surpass to receive a reward, was a function of performance success.

## RESULTS

We aimed to test the efficacy of both the joystick and the different automated training paradigms we had devised to teach a mouse to perform a reaching task. All of the water-deprived, adult mice were able to learn the task to criterion (>100 trials of >1cm reaches).

However, a considerable difference was noted between those animals that were exposed to the go cue, LIGHT EARLY rather than later on in training, the LIGHT LATE Condition (Figure 1). Unsurprisingly, the control animals in which no reaction time criterion existed performed best. Notably, within 30 sessions, animals were performing over 400 correct trials per session. ‘Correct’ was defined as a reach past the amplitude threshold and after the go-cue light came on. Conversely, ‘incorrect’ was defined as a reach past the amplitude threshold and before the go-cue light came on.

LIGHT EARLY animals leaned the reaction time task more quickly than LIGHT LATE, reaching criterion amplitude threshold roughly 8 sessions before LIGHT LATE on average (Figure 2). Additionally, once the punishment for reaching before the go cue was initiated, LIGHT EARLY mice had an overall lesser error rate, although the number of errors committed by each group were roughly equivalent.

**Figure 2:**
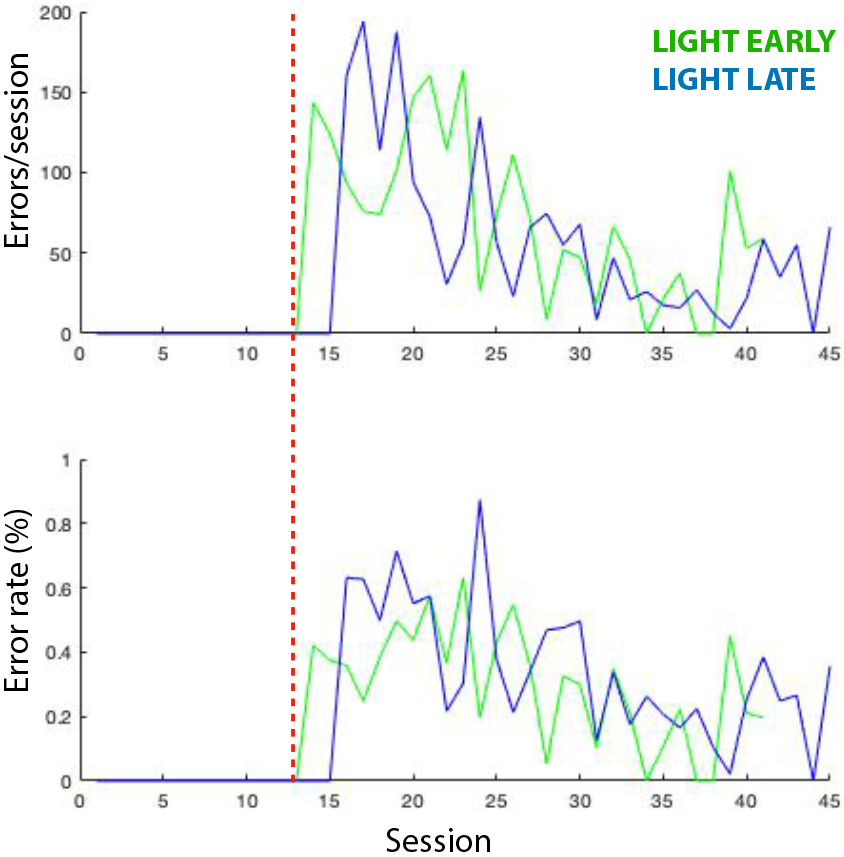
Errors committed as animals progress through either training regimen. Total errors (moving before reaction time light) and error rate (errors/total trials) for animals in either training regimen are shown top and bottom, respectively.

Finally, we observed that reaction time to the go-cue light reduced more quickly in the LIGHT EARLY training regimen. Reaction time was defined as the period of time between go-cue light on and surpassing the reach amplitude threshold. Median reaction time is shown in Figure 3. The average values are rather high due to our inclusion of the long rightward tail (All reaction times greater than 5 seconds were omitted). Even so, we can observe a steady decrease in reaction time, indicating that our mice learned the reaction time task. Further work will be performed to assess the generalizability of these observations to a larger cohort of animals and deepen our analysis of the highly-quantifiable data our mouse joystick provides.

## DISCUSSION

Studying the neural correlates of behavior requires precise, oftentimes real-time measures of those actions. In these early data, we demonstrate that 1) mice are capable of performing both a basic and reaction time reaching task and 2) introduction of the go-cue used in the reaction time task proved beneficial for learning the task. These results serve as a foundation for future work implementing tasks in mice that are normally thought to be reserved for primate research.

Scientific apparatus often come with a hefty price, precluding high-throughput use through multiple iterations of the device (Brunton et al., 2013) or purchasing any device to begin with. We document here an inexpensive, high-performance joystick to relay the position of reaches with sub-millisecond delay. As this is an open project, we encourage feedback and will update the methods and manners of implementation as our process matures.

**Figure 3:**
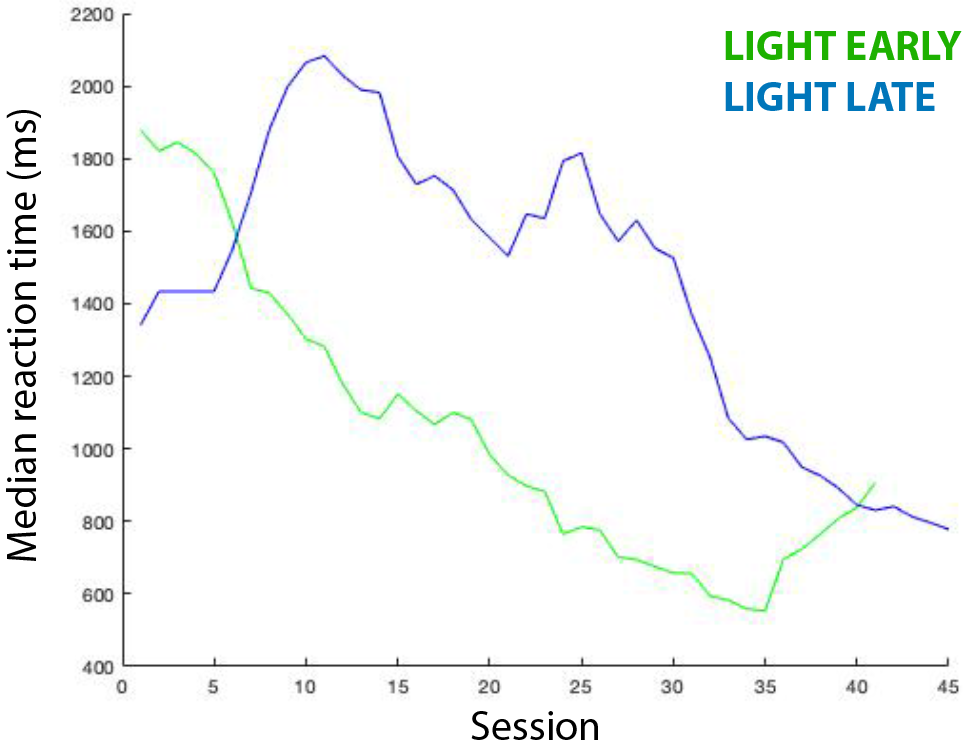
Median reaction times as animals progress through either regimen. Mice in the LIGHT EARLY task reduce their median reaction time more quickly. Median, rather than mean, was selected to minimize the effect of excessively long reaction times (mode was typically under 400ms, especially in latter sessions.

We describe here the pathway to build a head-fixed joystick setup – including hardware, data handling, and software. Also included are build and ordering instructions. This setup will work with any of the multiple mouse head-fixation solutions have been developed, including the RIVETS system used here (Osborne and Dudman, 2014). While already relatively inexpensive, we provide additional options to reduce cost, including using a 2-axis potentiometer joystick (~5$) in place of Hall effect joysticks (~75$). We prefer the latter, as the resistance is uniform in every direction - instead of having two axes along which there is less resistance. These tracks have the potential to skew the 2D trajectory of the reach, but this may not be a caveat for some experimental questions. Finally, if finer or more delicate reach kinematics are to be studied, the resistance of Hall effect joysticks may be decreased by cutting the spring – though care should be taken not to cut off too much (typically no more than 1.5 coils).

### Specific features of the joystick system described

- Arduino-based code for relaying multiple channels of data, including LED screen to display current task state and progress
- Rapid animal shuttle insertion, through magnetic platforms that instantly and reliably lock into a predetermined position
- Low cost, efficient build time, and small size can enable a lab to quickly and easily set up dozens of rigs in a small space.
- Task code to be run on rodents, including basic un-cued reaching, cued reaction time, and a probabilistic ‘two-arm bandit’ task requiring left or right reaches
- Analysis code for extracting reach performance parameters offline, including reach trajectory, amplitude, peak speed, duration, and inter-reach interval

## BUILD PROTOCOL: (see addendum at end of manuscript)

### TASK CODE

We have written code for several behavioral tasks common to non-human primate literature. Reaching has been utilized to study many important neural mechanisms, including the planning and generation of reaches, reaction time, and the valuation of different actions. As such, in our github repository (https://github.com/YttriLab/joystick) we have provided code for:

- un-cued reaching
- cued reaction time
- probabilistic reward ‘two-arm bandit’ task

### ANALYSIS CODE

We have produced data analysis code to quantify the execution of reaches, including reach trajectory, amplitude, peak speed, duration, and inter-reach interval. Detection is based on threshold crossing, and then works forward and backwards from this threshold to determine exact reach initiation and termination times. In doing so, the user is able to select for only full reaches and ignore small blips due to postural adjustment, grooming, or other non-task related behavior. For assistance with the code, we recommend contacting the lab directly in addition to visiting our github page, mentioned above.

## ACKNOWLEDGEMENTS

Andrew Buzza, Alex Hodge, and especially Joshua Dudman and his group at Janelia Research Campus.

---

**Materials List:** Headfixing Unit and Joystick Module

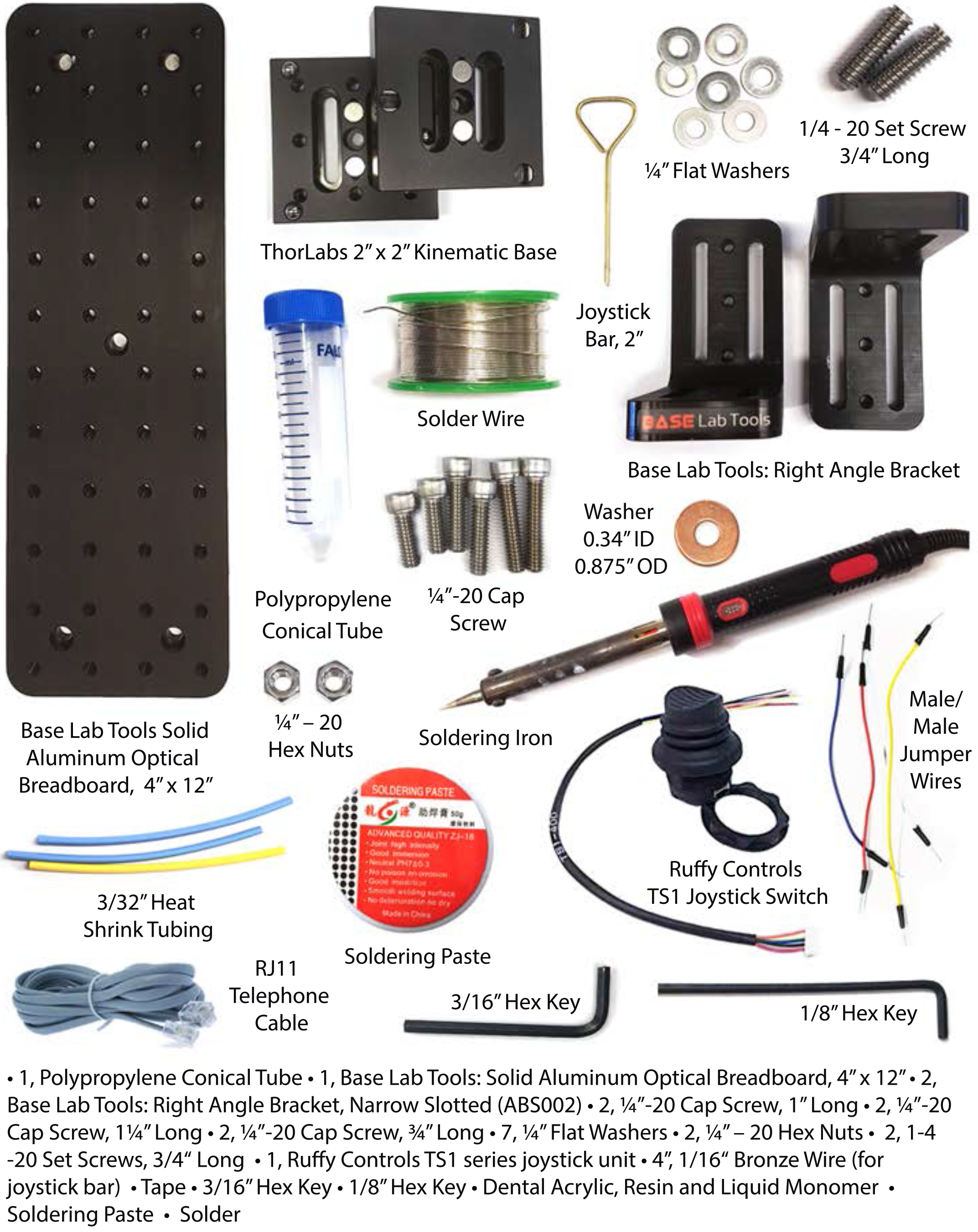

**Materials List:** Data Collection and Water Delivery

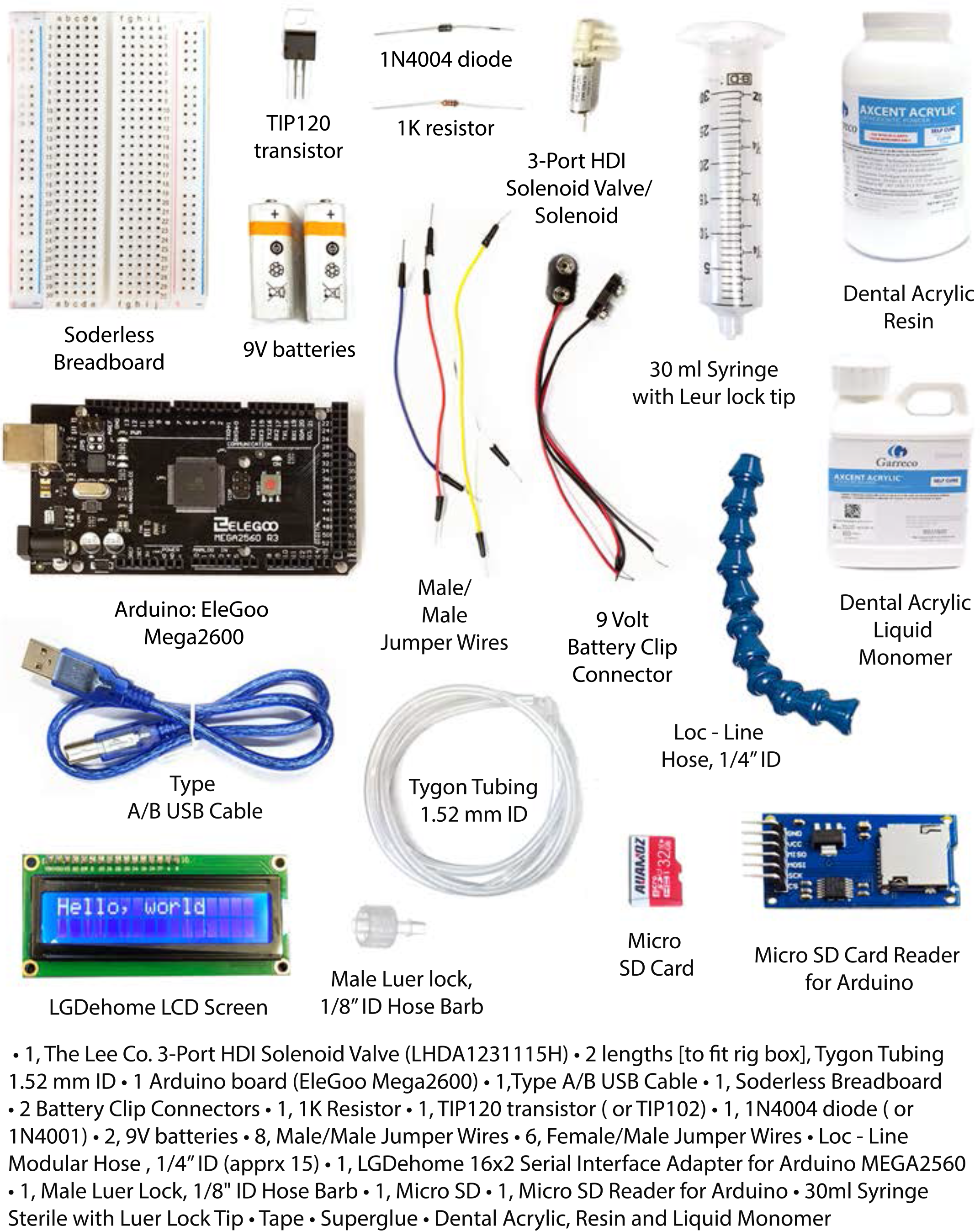

**Assembly:** Headfixing Unit

**Goal:** Build a headfixing device suitable for mice in electrophysiology experiments

1. Using AutoCAD files and instructions found on: **http://dudmanlab.org/html/rivets_designs.html** manufacture RIVETs stystem parts (forks, desired head caps) and shuttle components. Assemble into an adjustable shuttle with headfixing capabilities.

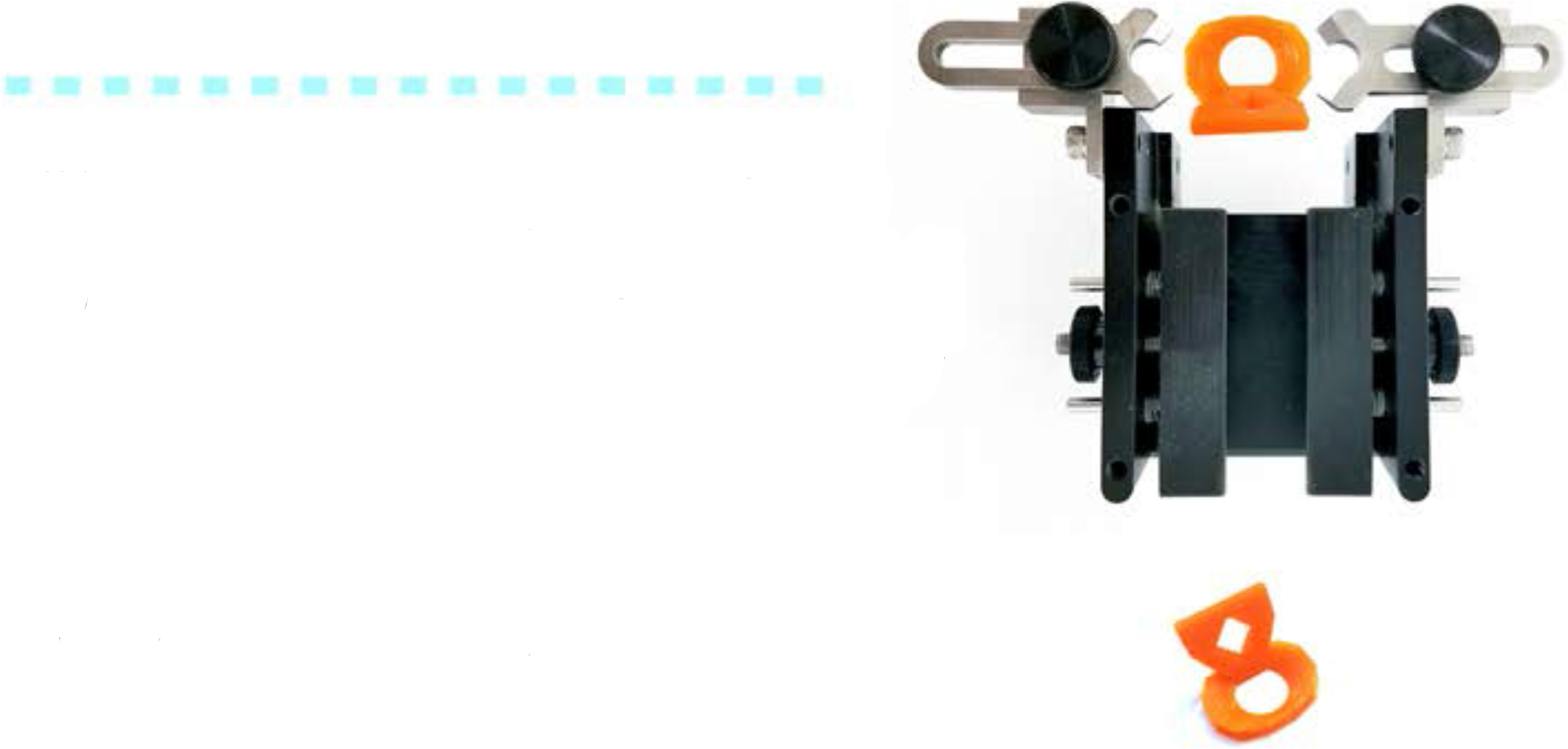

2. Based on the Methods section of: **Osborne & Dudman (2014) PloS One 9(2): e89007** permanently attach head caps to experimental mice.

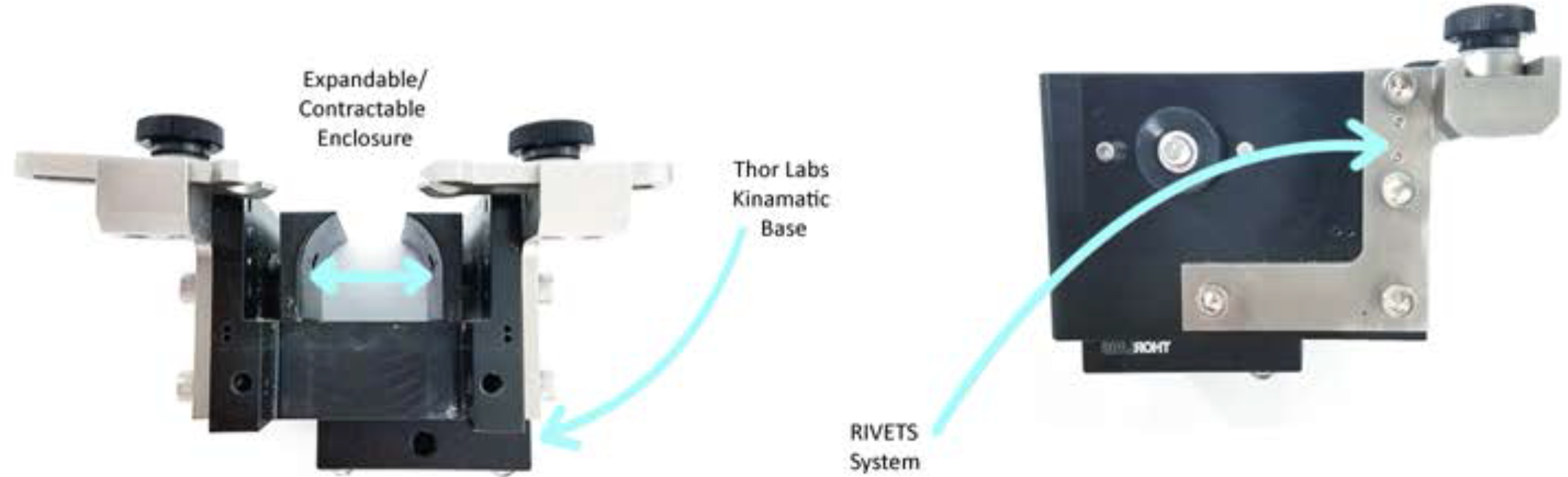

**Assembly:** Headfixing Unit

**Goal:** Build base and stand for headfixing shuttle. Steps:

1. Hold 2 right angle brackets in a ″Z″ shape, so that the slots align. Thread a 11/4″1ong, 1/4″ - 20 cap screw with a 1/4″ f1at washer, and push through both bracket slots.
2. Cap the emerging end of the screw with a 1/4″ flat washer, and then a 1/4″ - 20 hex nut in order to hold the 2 brackets together. Join the pieces at the lowest extension level possible so adjustments can be made in later steps.

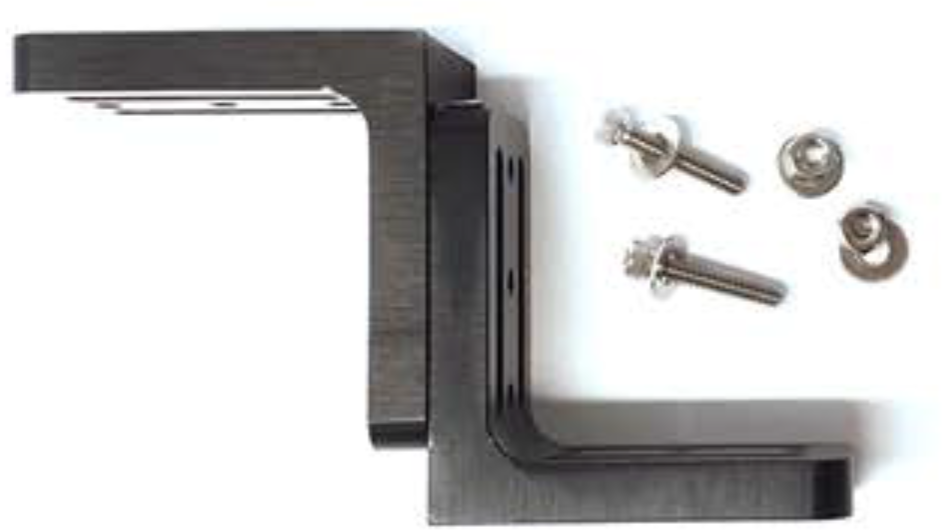

3. Repeat steps 1 and 2 with the other slot for security. In order to fully tighten hex nuts onto the screws hold the cap of the scew in place with a 3/16″ hex key, and tighten the nut with a wrench or pliers.

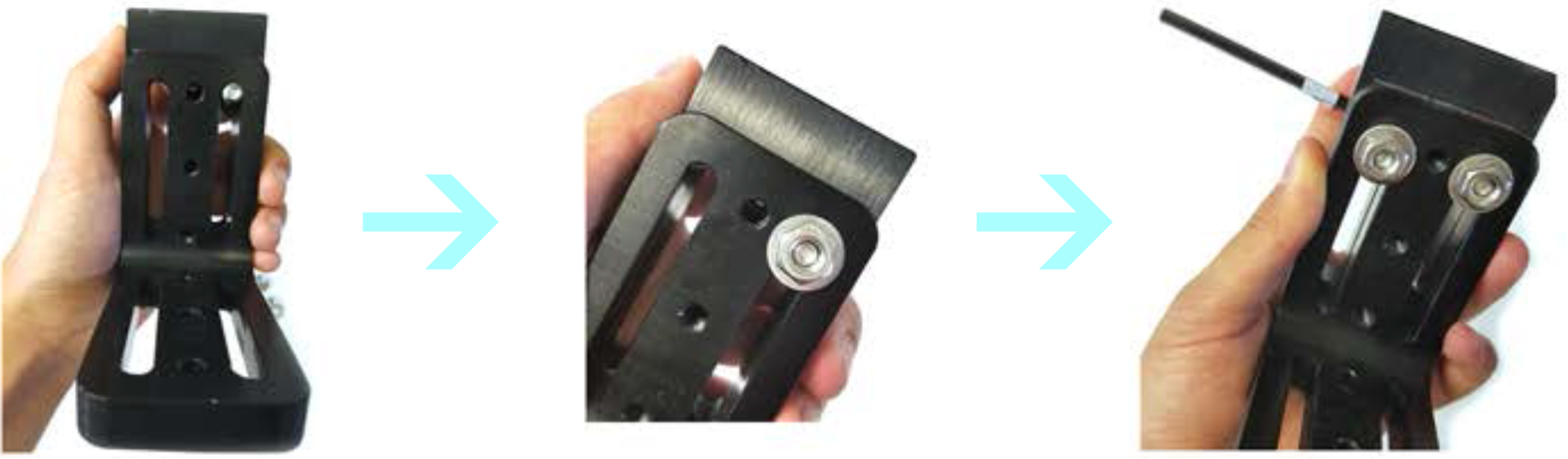

4. Screw the bracket unit into the optical breadboard with a 5/8″ long, 1/4″ - 20 cap screw threaded with a 1/4″flat washer. The unit should be placed to lower right handside (not centered), in order to account for offset of the headfixing shuttle. Only use one screw to hold the bracket unit to the breadboard so the unit can be rotated.
5. The kinematic base has 2 sides -- one with protruding balls and the other with wells. Screw the side with 2 wells into the screw space closest to the top of the board with a 3/4″ long, 1/4″ - 20 cap screw.

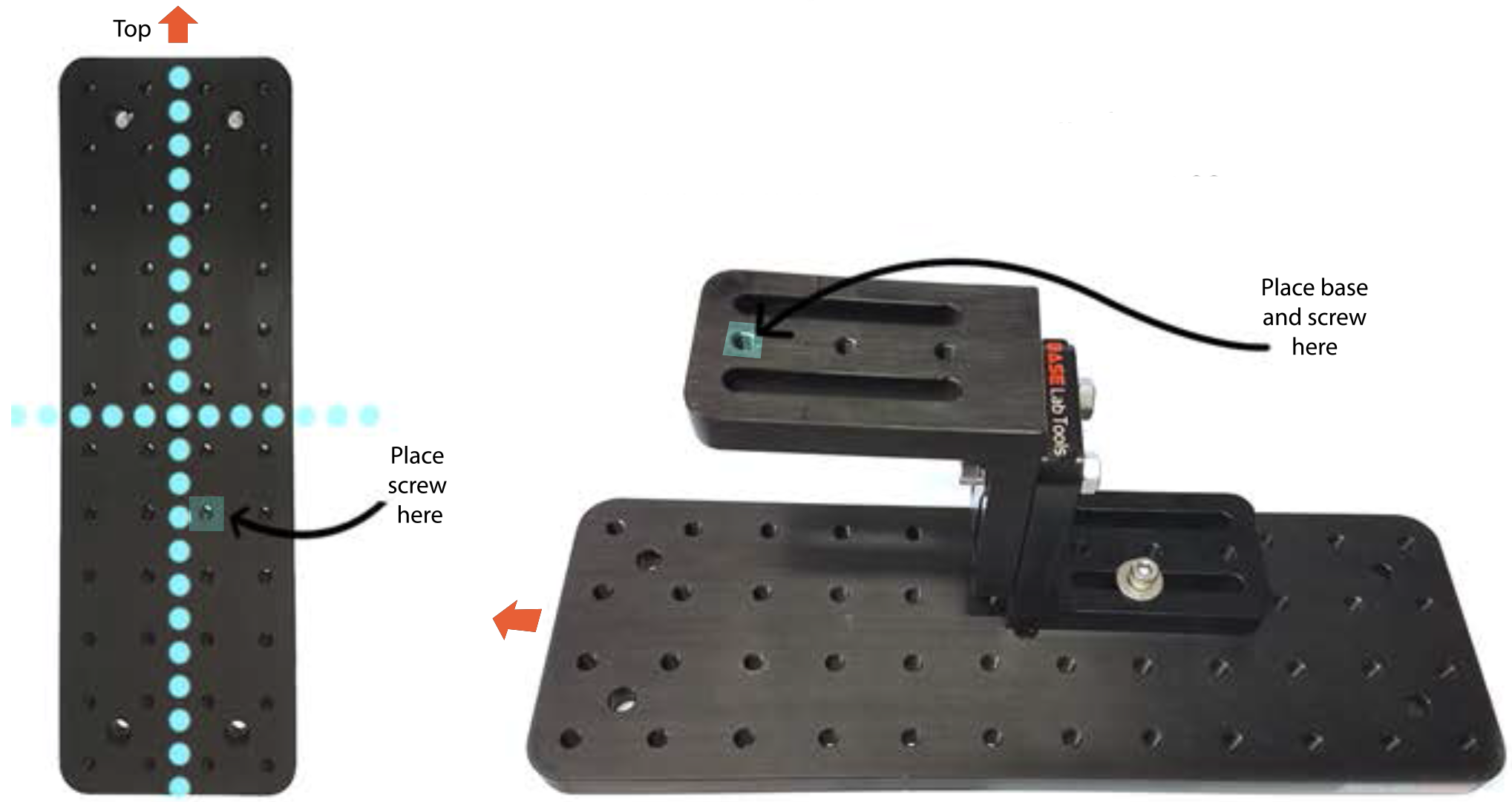

**Assembly:** Joystick Stand

6. Screw the side with balls to the headfixing shuttle by the top right corner of the base with a set screw with the 1/8″ hex key.

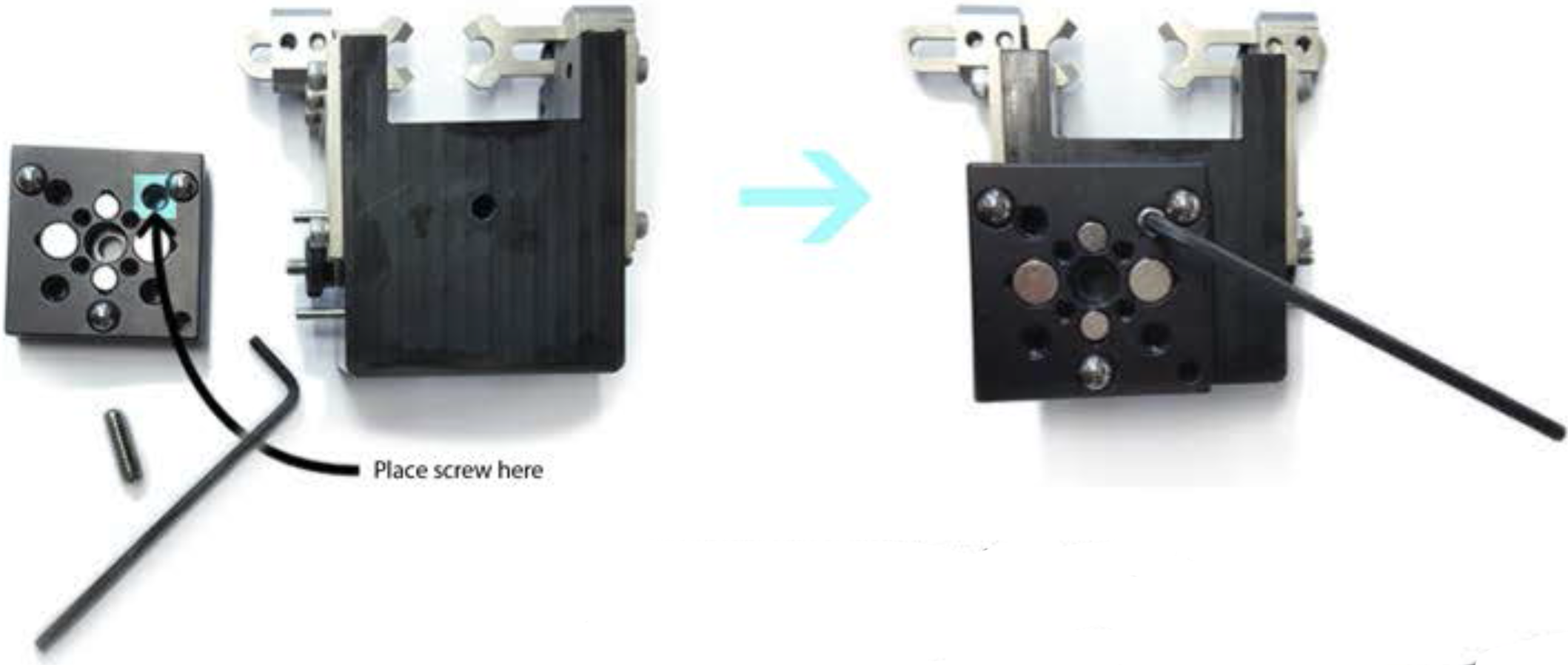

7. Mark off a 50 ml plastic conical tube at 10 ml. Using a dremel or other plastic safe cutting tool, carefully drill a hole slightly larger than a 1/4″ - 20, 5/8″ long screw into the still attached cap. If using a dremel: Unscrew the cap from the tube, and screw upside down on the breadboard with a washer, so that the threads of the cap are pointed up. Screw the tube into place, to make a sort of stand for trimming off the pointed end of the tube at the 10 ml mark. Cut the tube by drilling into the tube, and dragging the inserted tip around the circumfrence of the tube, ensuring that the sides remain even.

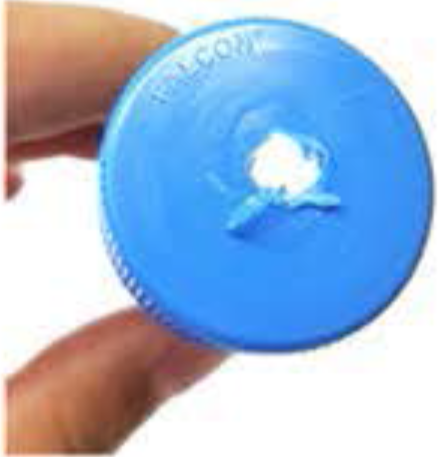

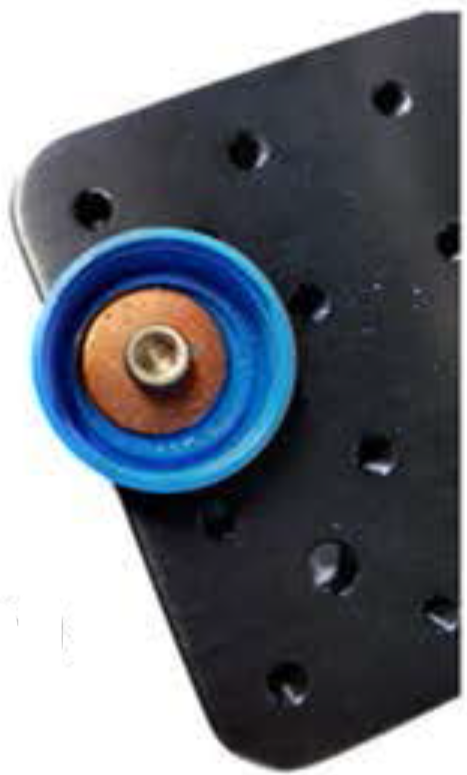

8. Drill a dime sized hole in the tube, near the base of the joystick unit so the wires can be threaded through the tube. Remove wires from joystick base, and unscrew cap from the board.
8. To make Joystickbar, snip 1/16″ bronze wire into 4 inch segments. Bend the wire at a 90° angle an inch from the end to form a handle.

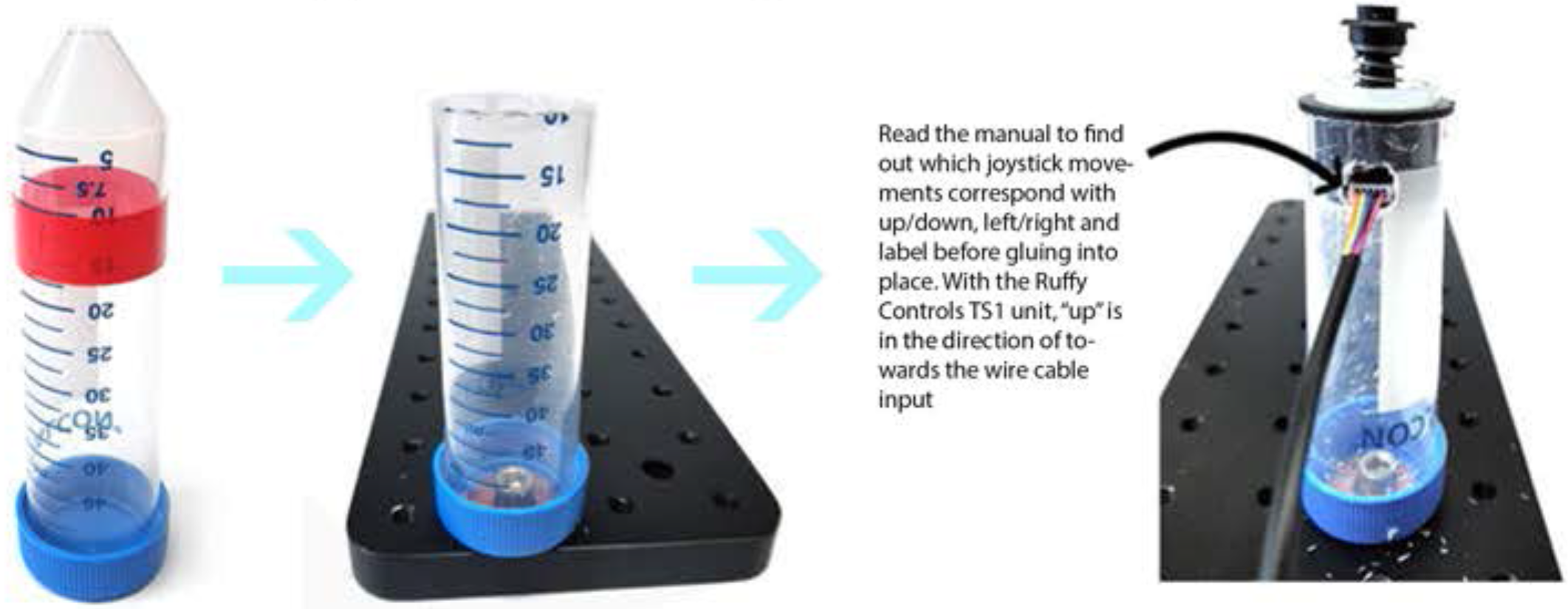

**Assembly:** Joystick Stand

9. Tape around the bottom of the cap to prevent leaks. Place a small amount of dental acrylic between the washer and cap. Screw the cap and washer into the board to act as a clasp, and fill the bottom of the cap with dental acrylic, avoiding the screw head and threading of the cap so the rest of the tube can still be screwed in. Allow to dry completely.

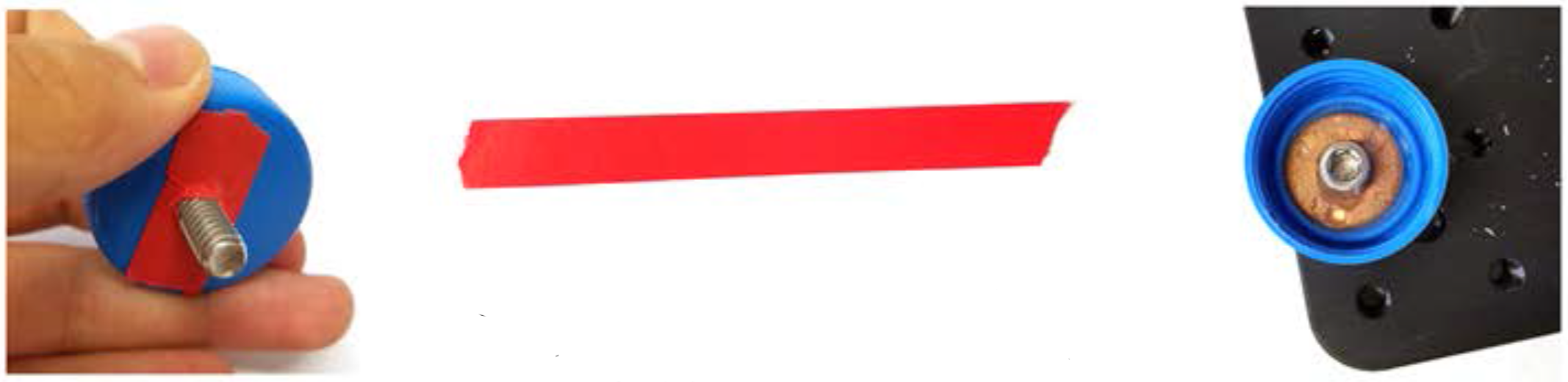

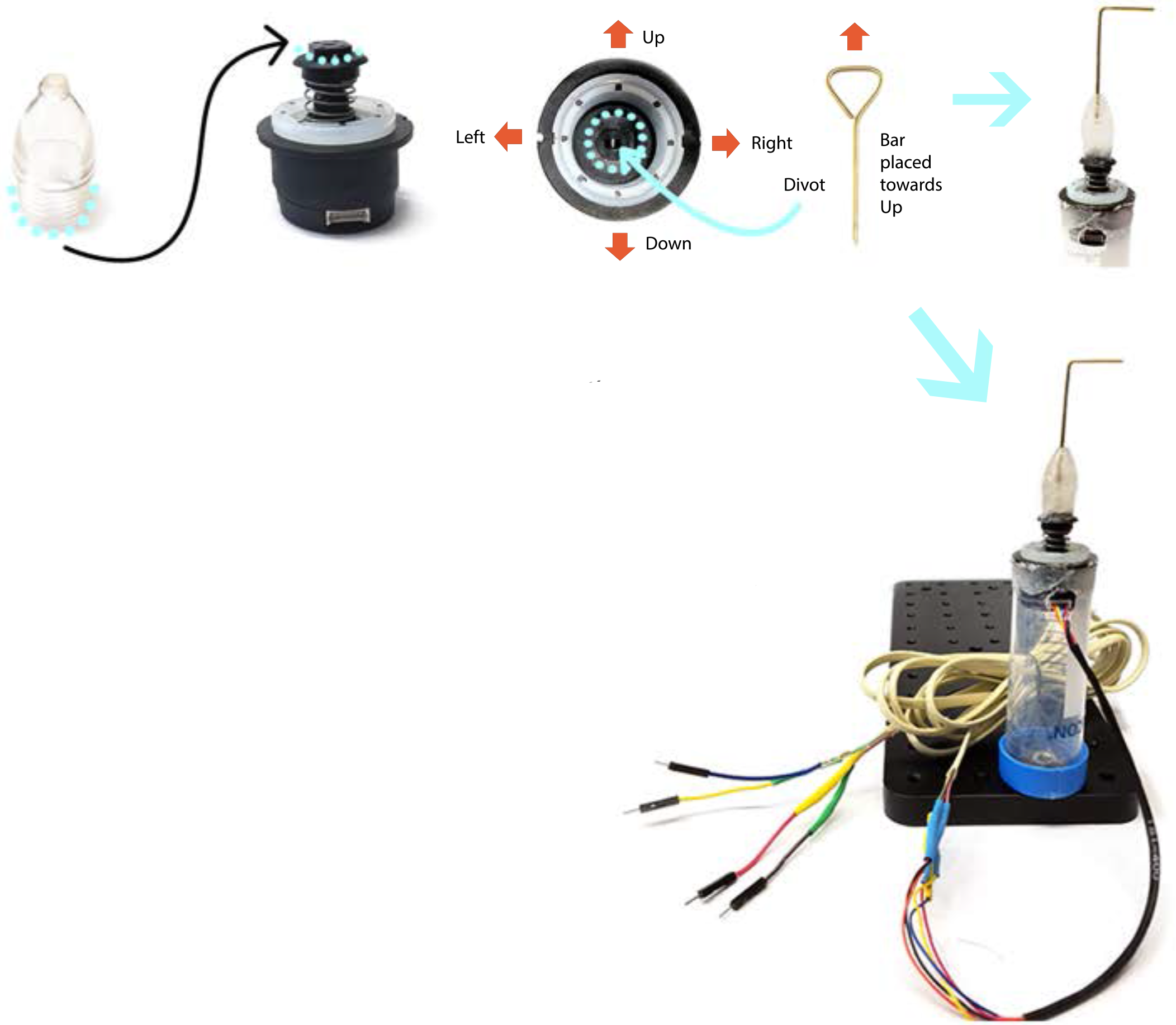

10. Remove the rubber cap from the top of the joystick unit. In order to attach the bar to the joystick unit, fill a cap that fits over joystick head with dental acrylic, and push it over the top, so that the dental acrylic settles on to the base. Push the joystick bar into the cap through the dental acrylic, and into the small central divot in the top of the joystick base. Make sure the bar and handle are straight, and that the handle is aligned with being pushed up and down. Allow to dry completely.
11. To attach the unit to the stand, smear a stiff dental acrylic mixture around the base of the unit, and the top of the stand, avoiding the hole meant for joystick connections. Push into place, while lining up the wire input and exit on the joystick and the stand. Allow to dry completely.
12. Once all pieces are dry, reassemble. Screw the base in so that the stand and joystick have ″Up″ pointing forward. Mark on the tube and cap what forward. Mark on the tube and cap what forward is, so if the stand needs to be disassembled it can be reassembled wiht ease.

**Assembly:** Data Collection

**Goal:** Attach joystick wires to cable wires in order to extend joystick communication to the Arduino board.

1. Strip about an inch of plastic coating from the ends of joystick wires and 4 male end jumper cables with wire strippers to expose inner metal threads. Cut away about 2 inches of plastic casing from both ends of an RJ11 cable to reveal coated wires, and strip plastic away. Take care to not break too many metal threads in the stripping process, to ensure a strong electrical connection once the soldering process is complete.

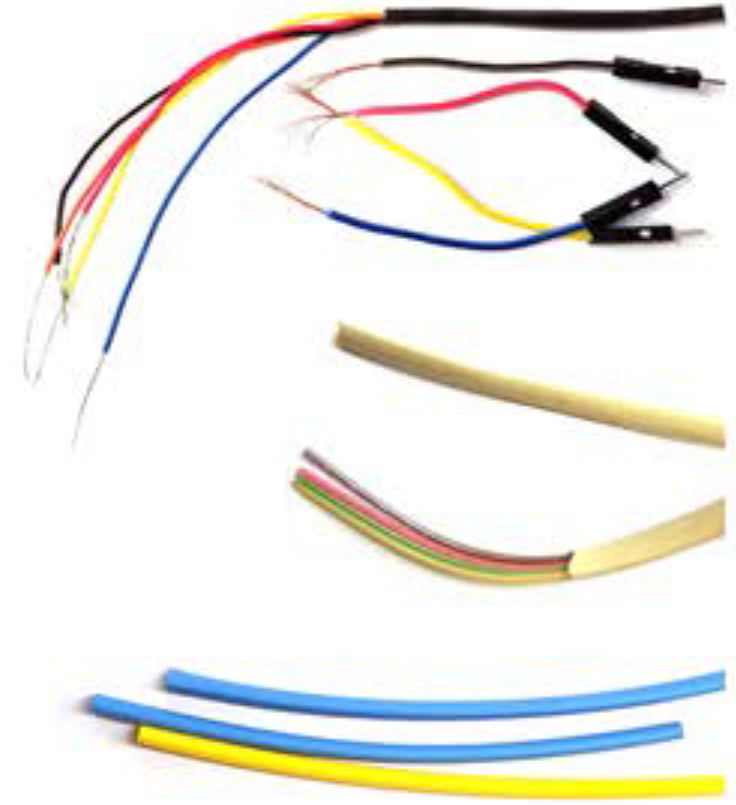

2. Check the joystick manual to see what color wire corresponds to what information sent. If certain Information is uneeded (ex: not collectiong ″push button″ data), plan to solder that wire with the joystick ″ground″ wire to the RJ11 cable, because the joystick wlll not work if any wires are not plugged in to the Arduino board.
3. Heat up soldering iron, and cut shrink wrap tubing into 8, 1″ pleces. With ″helping hands″ or another clamp system that can hold wires in proximity to each other, arrange a joystick wire across from a slmllarly colored wire in the cable (for clrity -- RJ11 cable colors do not have particular meanings like the joystick wires do). To use soldering iron: Once solder wire melts when touching iron, it is hot enough to hold connections between separate wires. Dip tip of iron in soldering paste, and hold it to the exposed wire to heat. With the iron stlll touching the wire from behind, touch solder wire to the from right above the desired surface, so that as it the solder wire melts, it flows on to and coats the appropriate surface.

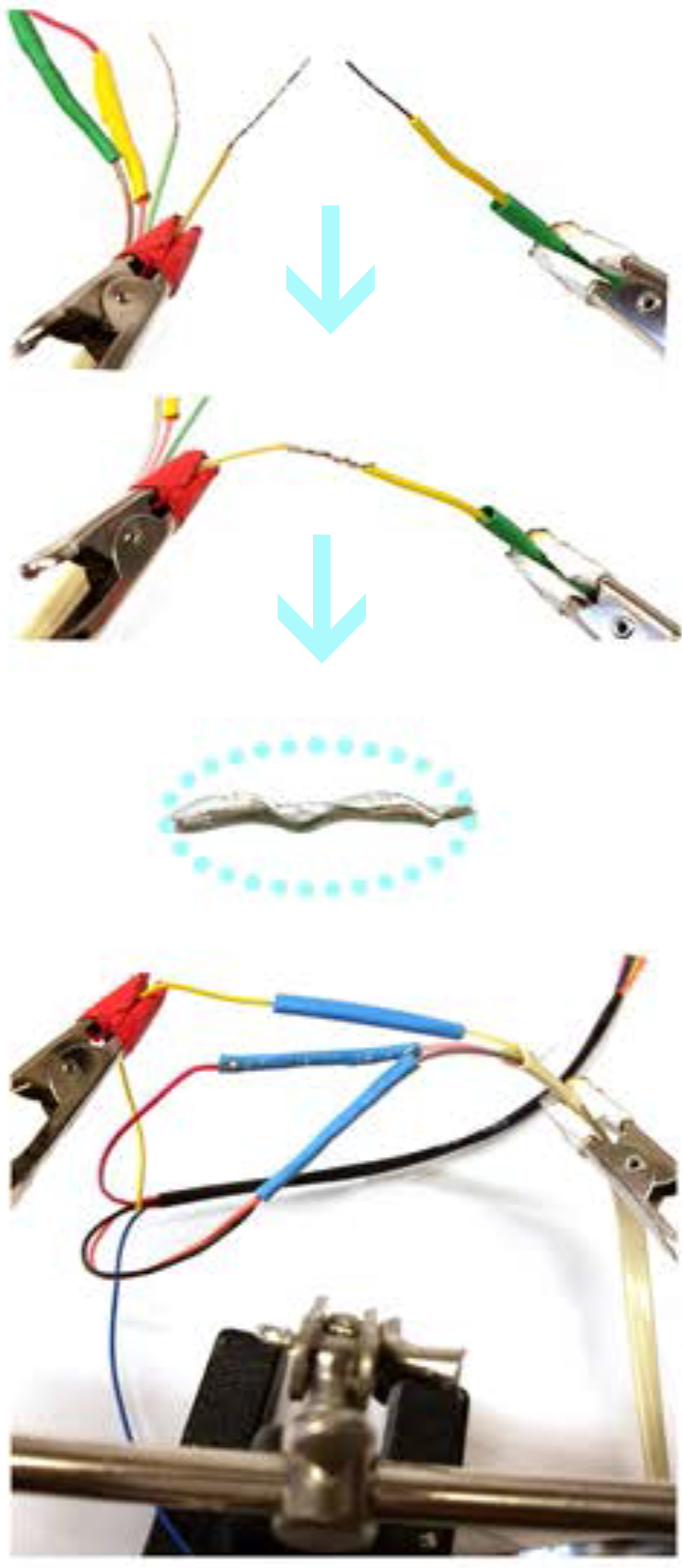

4. Thread a section of shrink wrap tubing over one of the wires. ″Tin″ both lengths of exposed wire by coating stripped area with a thin layer of solder. Make a mechanical conection between the 2 wires by intertwining, and secure the connection by melting a thin layer of solder over the surface, until no gaps remain. If the solder balls up, there is too much on the connection. Reheat the area until solder is liquid, and remove excess.

**Assembly: Data Collection and Water Delivery**

6. Slide the shrink wrap tubing over the junction, so that wires are not exposed. Heat the shrink tubing with the side of the unplugged soldering iron to form the plastic to the connection.
7. Repeat soldering and shrink wrap process for all other joystick wires, remembering to join uneccessary wires together with ground.
8. On the other end ofthe RJ11 cable, solder male ends to corresponding colors so the cable can more easily be plugged into the Arduino board.

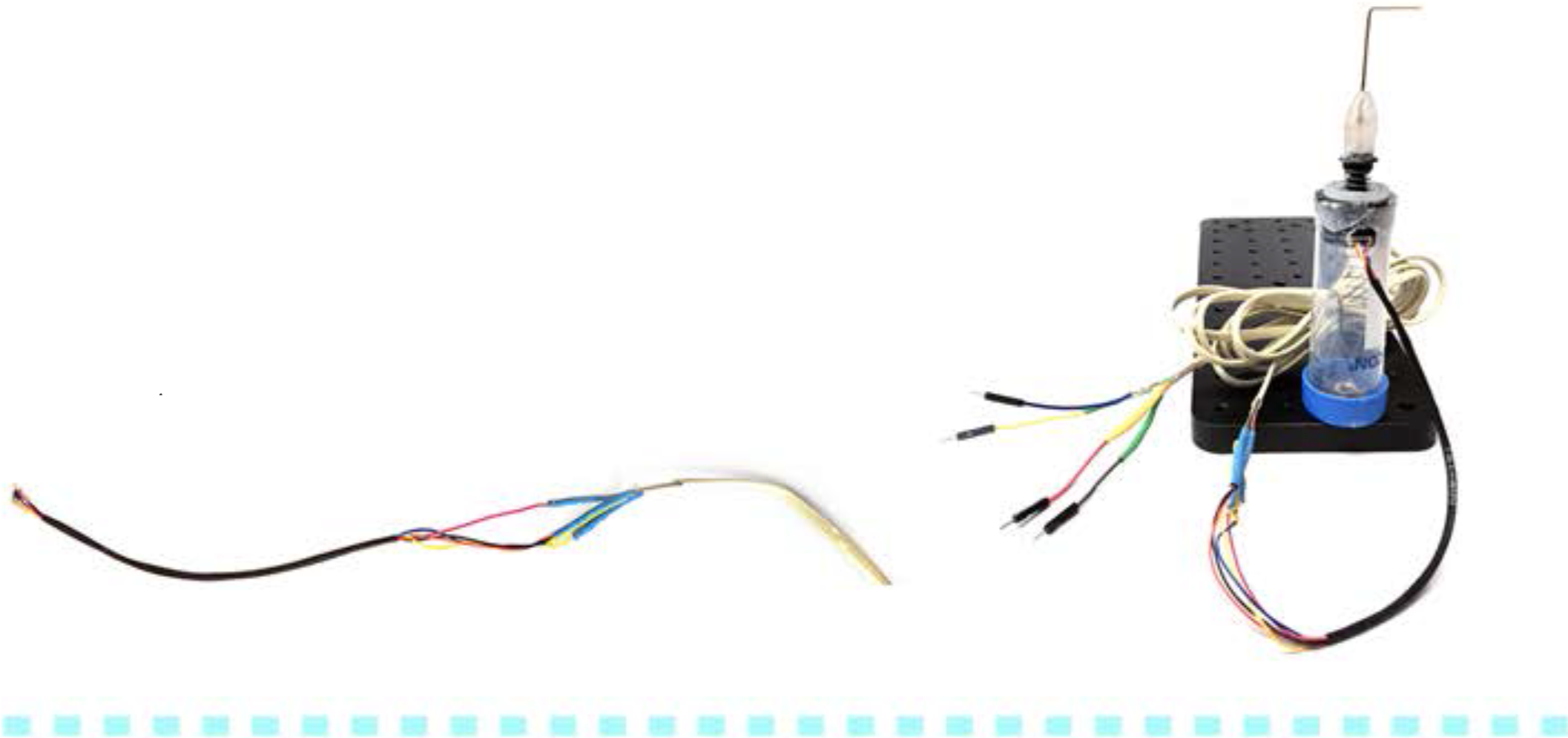

**Goal:** Attach screw to Loc-Line hose so that the hose can be screwed into the optical breadboard to scaffold deliver water tubing

1. Mix equal parts of dental acrylic resin and liquid monomer in a disposable dish, wearing gloves. Consistency will be runny, and stiffen until completely hard (~ 10 minutes).

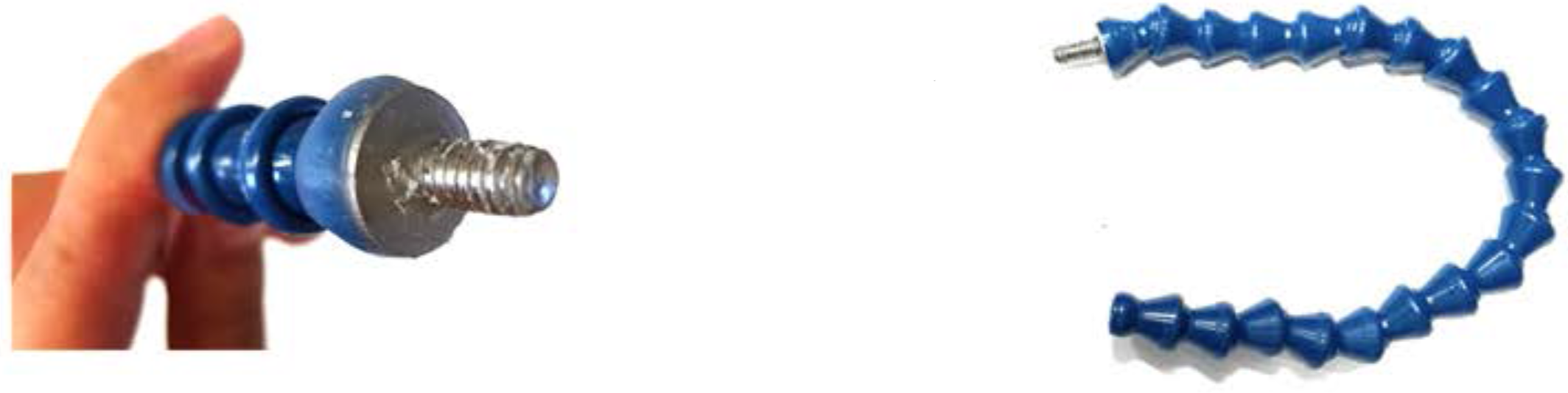

2. Snap together a line of approximately 15 Loc-Line pieces. Fill the base (flared end) of the last piece with dental acrylic mixture, taking care to coat sides and fill center. Embed cap screw head in the mixture, so that it is centered in the base and is surrounded by dental acrylic.
3. Push the washer Over the screw, until it is aligned with the bottom of the base Loc-Line piece. Cover the washer and outside ofthe base with dental acrylic and mold mixture around both in order to secure the screw and washer in place.
4. When the dental acrylic has is mostly stiff, make sure screw is in a straight line with the rest of the hose and that dental acrylic is not in the threads of the screw. Hang to dry with the screw pointing down so that dental acrylic inside the base piece settles around the screw head. Allow to dry completely overnight.

**Assembly:** Water Delivery

5. Screw Loc-Line hose into the top left corner of the optical board. Make sure the head fixing unit and the joystick stand are aligned so the joystick bar is underneath the forks of the shuttle.
6. Affix water delivery port of choice to Loc-Line hose.

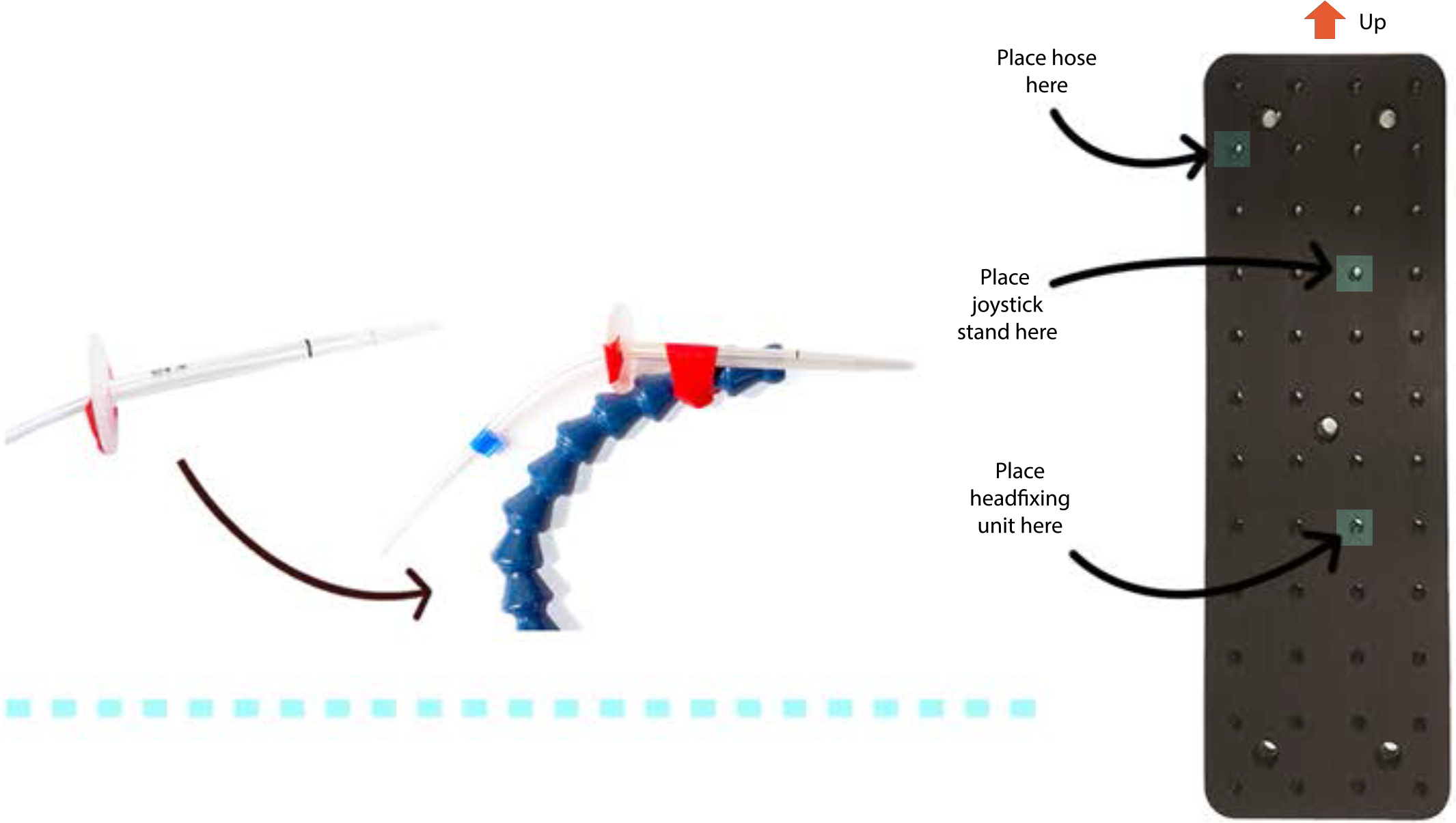

**Goal:** Build an Arduino controlled water delivery system and connect to joystick and headfixing unit

1. Using onllne Instructions. create an Arduino controlled solenold. Instructions found at: **https://www.instructables.com/id/Controlling-solenoids-with-arduino/** Look at the schematic wiring diagram to check that current is flowing through the entire circuit (ex: resistor and diodes are pointing the correct way), and that the solenoid is properly connected.

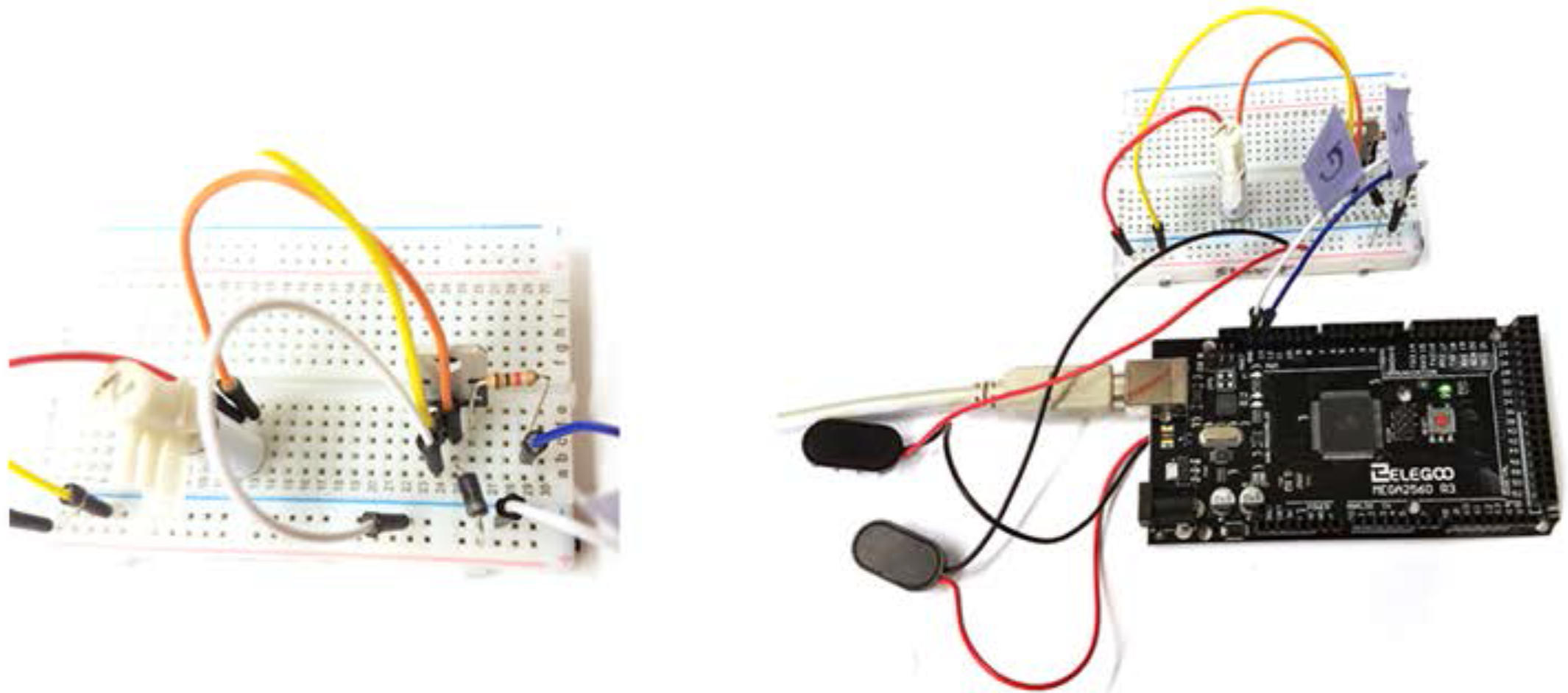

**Assembly:** Data Collection and Water Delivery

2. Once circuit is complete, attach lick spout tubing to the output value of the solenoid. Attch other length of Tygon tubing to the input value. If using the 3-Port HDI solenoid, leave the third valve capped.

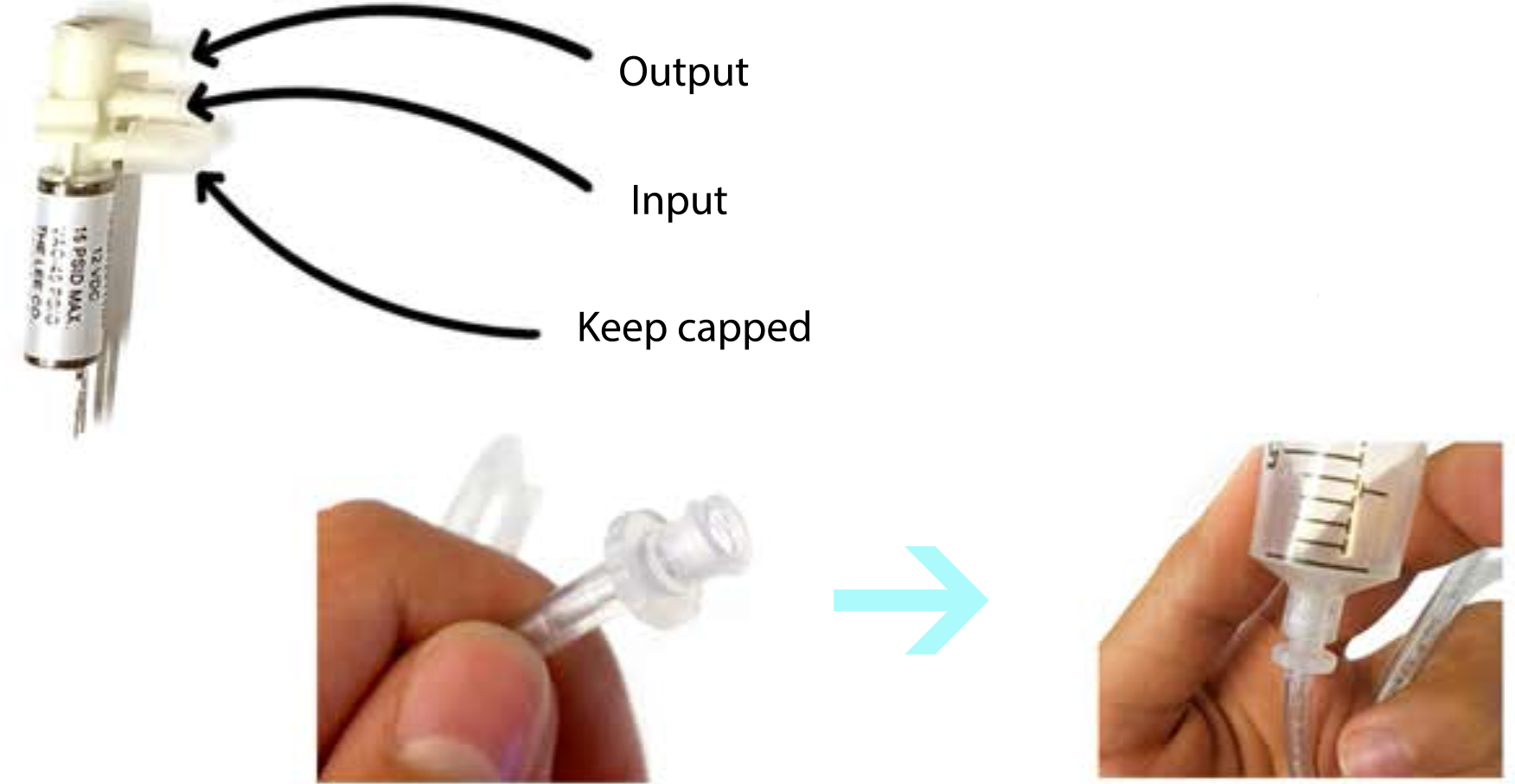
3. Push unconnected end of input valve tubing over Luer loc screw, and screw into base of Luer lock 30ml syringe.
4. Connect LCD screen to the Arduino board, so that mouse progress can be tracked in real time. Using the back of the screen as a guide, connect the screen to the Arduino board through ports SCL, SDA, Ground, and 5V with four male/male jumper cables.
5. For final assembly of system, tape tube connected syringe on to a wall near where the rig will be kept. Arrange Arduino board and electrical breadboard onto optical breadboard with headfixing unit, to create a contained, modular training system. Secure down pieces with tape or twist ties.

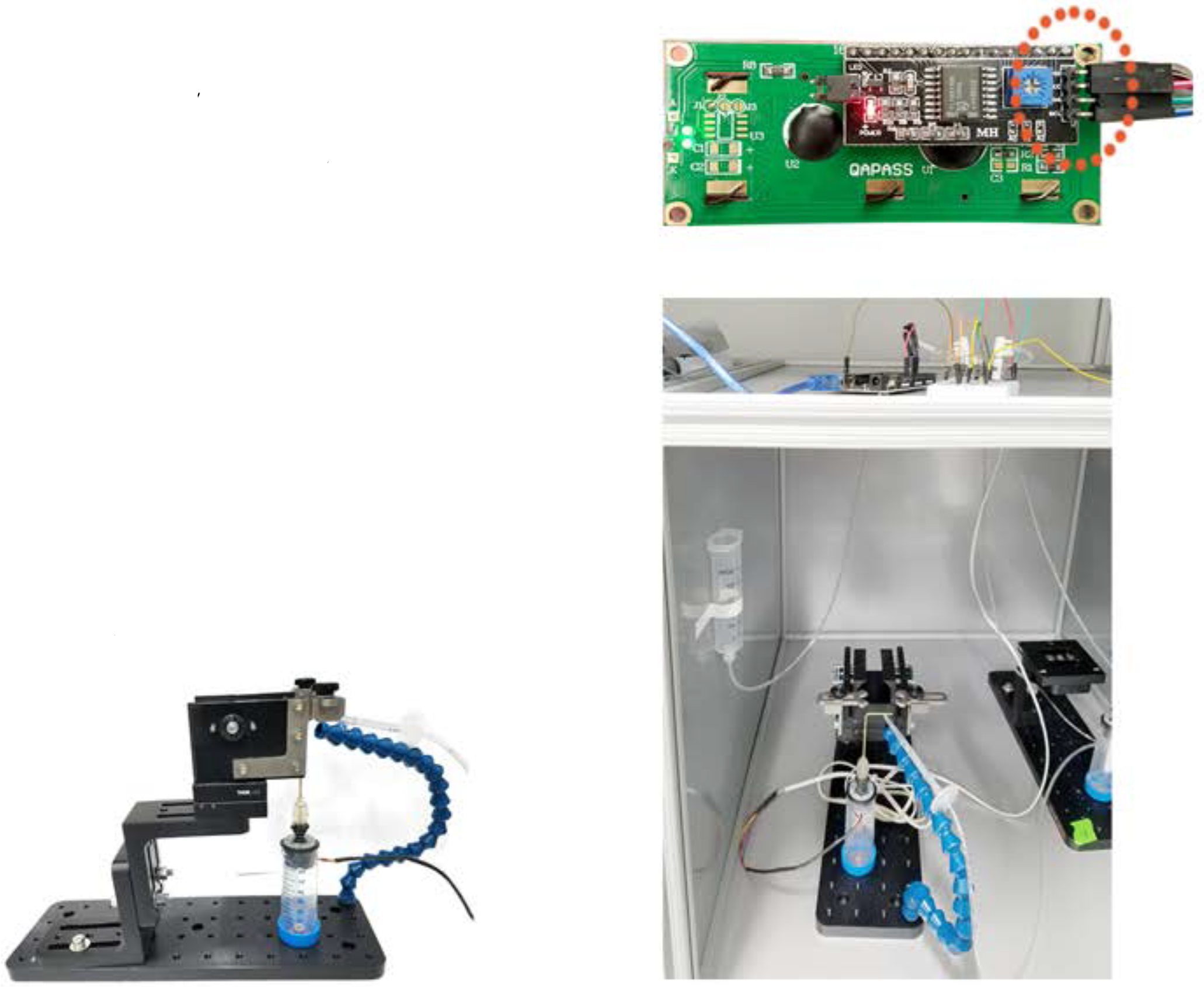

## Notes on Data Collection

Because neither the Arduino IDE nor board store data, joystick information must be held in alternate programs like Processing, or on additional hardware, like micro SD cards. Through both collection methods, .csv files containing desired information can be generated and read through data processing software such as MATLAB.

### Processing

1. Set up a Processing script to write files to a certain folder path.
2. Make sure board is directly connected to computer, by way of the A/B USB cable.
3. Select the correct port in Arduino, from the pull down menu path: Tools>Port
4. To record data, run the Processing script (play button). Check that the file is actively writing to the desired folder. If it is not, stop the script, unplug/re-plug the A/B cable, and try again.

### Micro SD

1. Set up MATLAB script to pull files off of micro SD card.
2. Attach card reader to Arduino board, based on inputs (CS, SCK, MOSI, MISO, VCC, GND) on the back of the device with 6 Female/Male jumper cables.
3. When recording data, make sure micro SD card has “clicked” into place in the reader. To stop recording, take the SD card out of the reader.

### Software links

Arduino IDE – https://www.arduino.cc/en/main/software

Processing – https://processing.org/download/

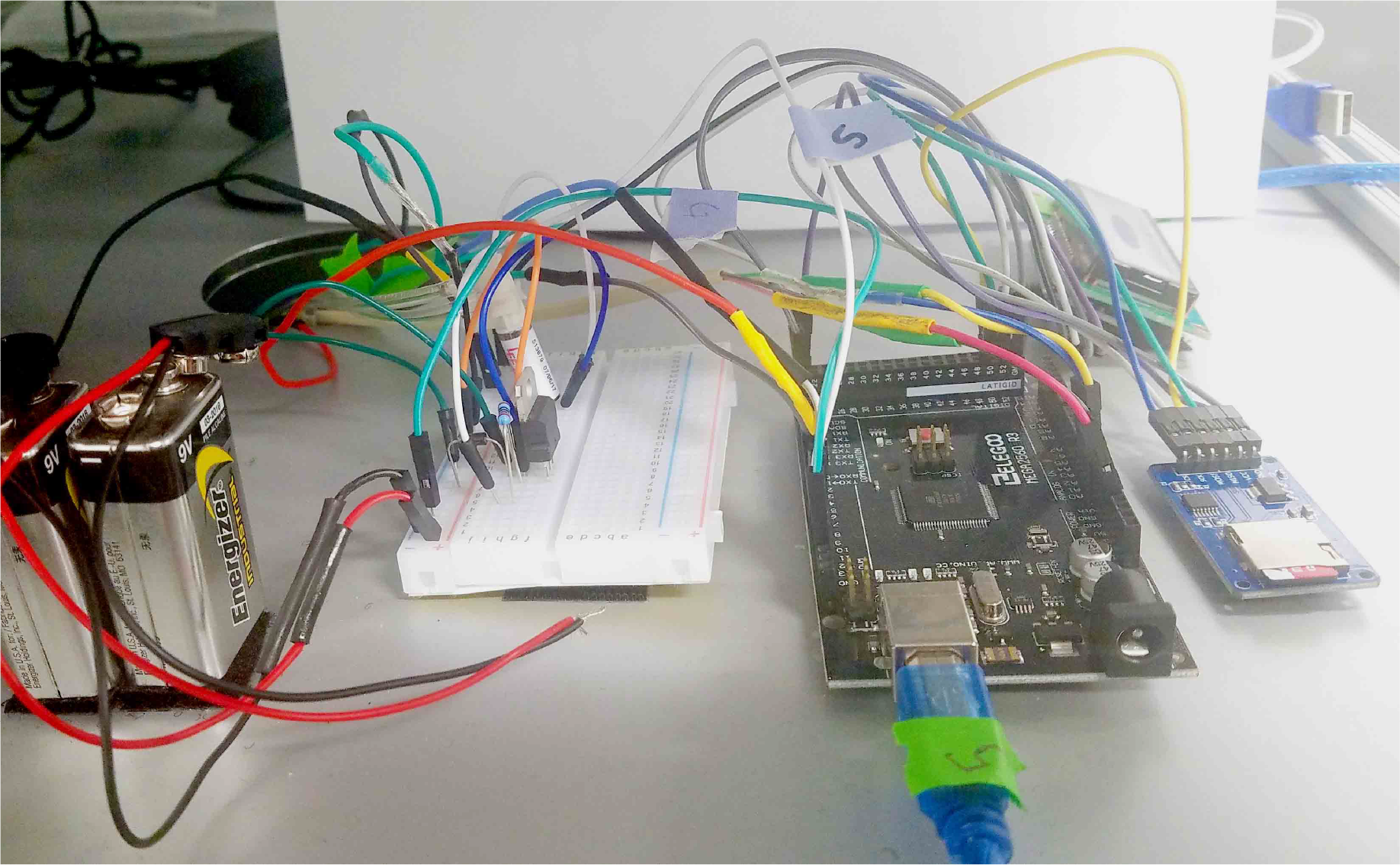

## Materials Order Form

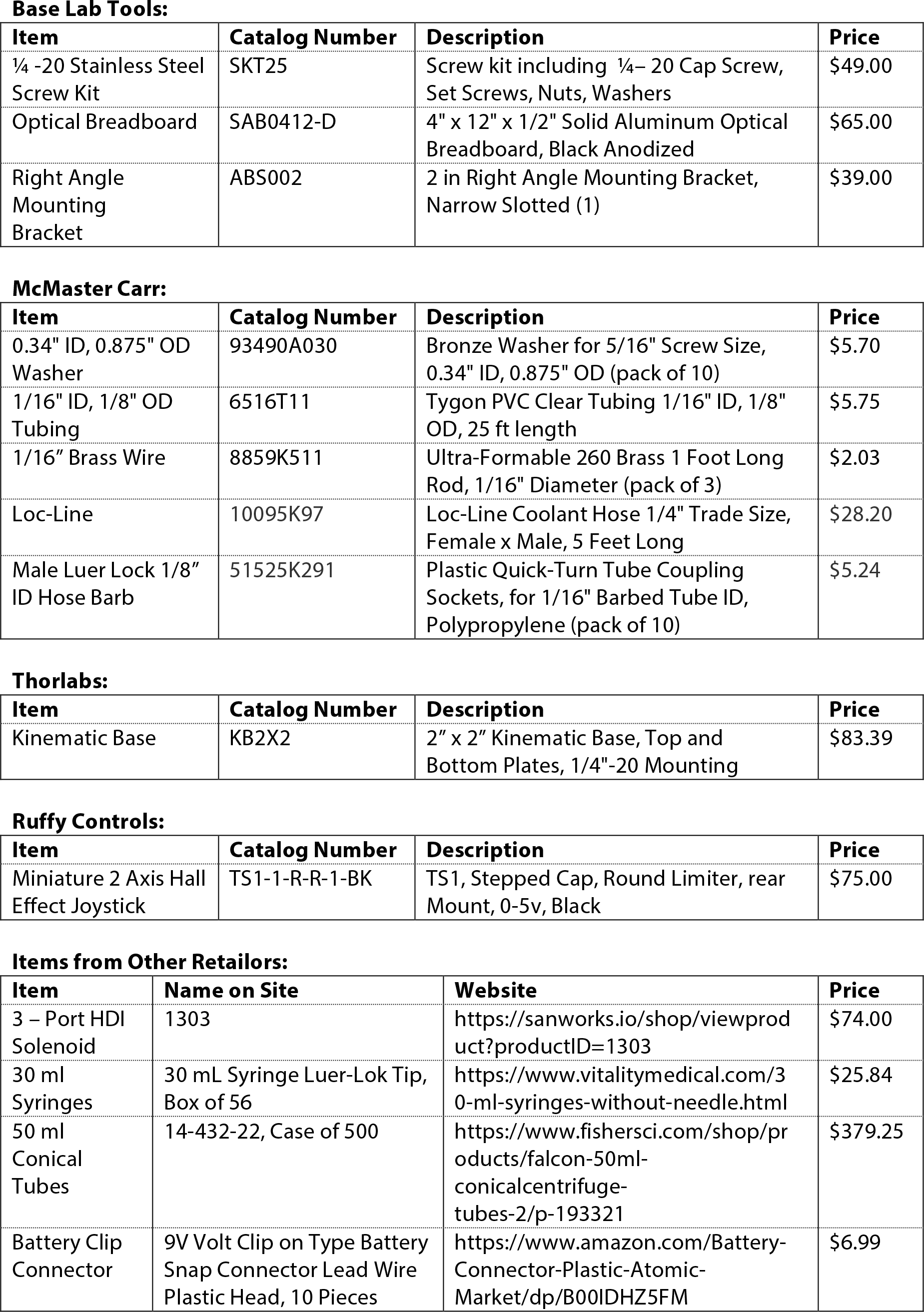

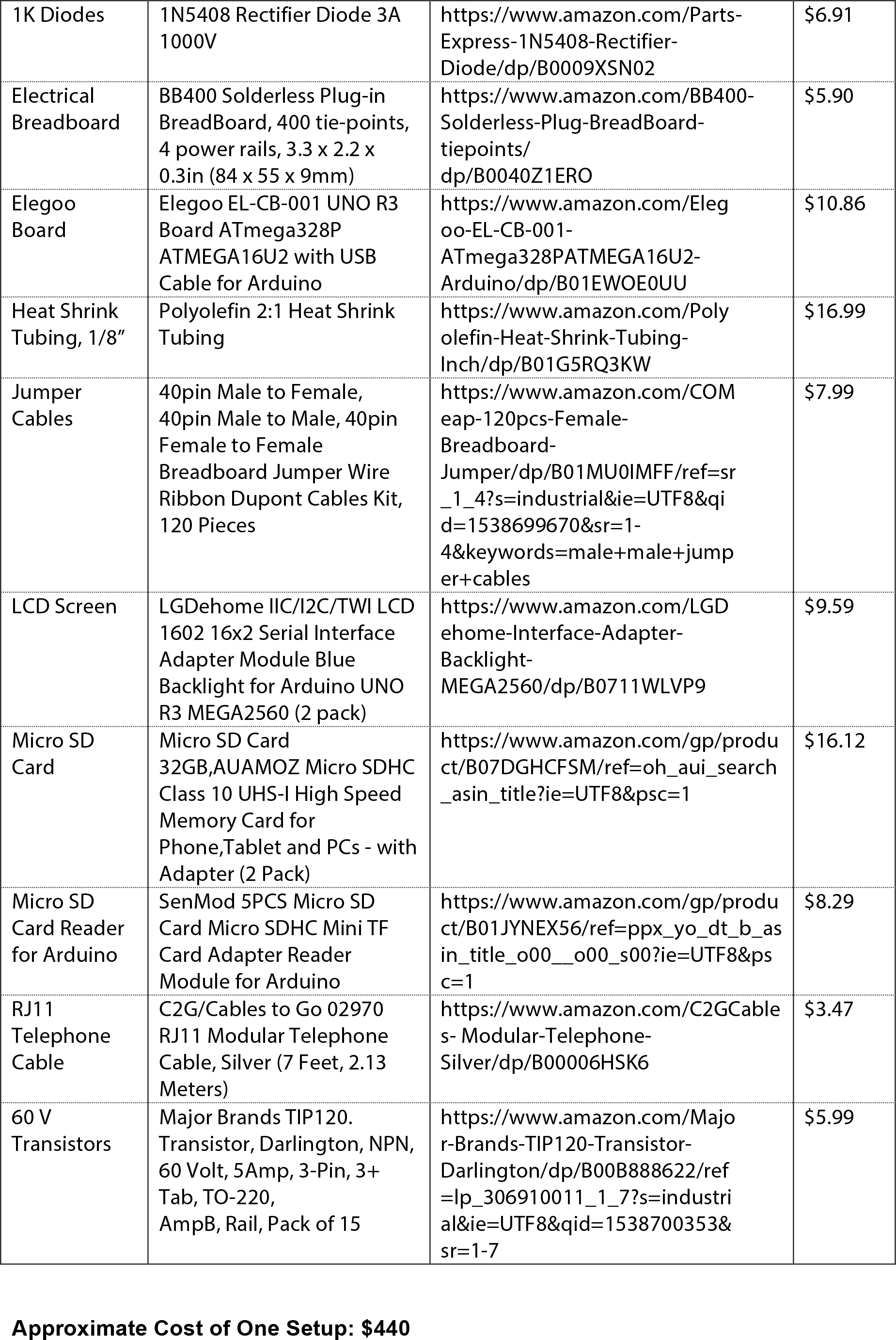

## REFERENCES

Bollu, T., Whitehead, S.C., Prasad, N., Walker, J.R., Shyamkumar, N., Subramaniam, R., Kardon, B.M., Cohen, I., Goldberg, J.H., 2018. Cortical control of kinematic primitives in mice performing a hold-still-center-out reach task. bioRxiv 304907. https://doi.org/10.1101/304907

Brunton, B.W., Botvinick, M.M., Brody, C.D., 2013. Rats and humans can optimally accumulate evidence for decision-making. Science (80-.). https://doi.org/10.1126/science.1233912

Cherian, A., Fernandes, H.L., Miller, L.E., 2013. Primary motor cortical discharge during force field adaptation reflects muscle-like dynamics. J. Neurophysiol. 110, 768–783. https://doi.org/10.1152/jn.00109.2012

Churchland, M.M., Cunningham, J.P., Kaufman, M.T., Foster, J.D., Nuyujukian, P., Ryu, S.I., Shenoy, K. V., Shenoy, K. V., 2012. Neural population dynamics during reaching. Nature. https://doi.org/10.1038/nature11129

Cohen, J.Y., Amoroso, M.W., Uchida, N., 2015. Serotonergic neurons signal reward and punishment on multiple timescales. Elife 1–25. https://doi.org/10.7554/eLife.06346

Dean, H.L., Hagan, M.A., Pesaran, B., 2012. Only Coherent Spiking in Posterior Parietal Cortex Coordinates Looking and Reaching. Neuron 73, 829–841. https://doi.org/10.1016/j.neuron.2011.12.035

Ellens, D.J., Gaidica, M., Toader, A., Peng, S., Shue, S., John, T., Bova, A., Leventhal, D.K., 2016. An automated rat single pellet reaching system with high-speed video capture. J. Neurosci. Methods. https://doi.org/10.1016/j.jneumeth.2016.07.009

Fetsch, C.R., 2016. The importance of task design and behavioral control for understanding the neural basis of cognitive functions. Curr. Opin. Neurobiol. 37, 16–22. https://doi.org/10.1016/j.conb.2015.12.002

Fromm, C., Evarts, E. V, 1981. Relation of size and activity of motor cortex pyramidal tract neurons during skilled movements in the monkey. J. Neurosci. 1, 453–460.

Guo, J.-Z., Graves, A.R., Guo, W.W., Zheng, J., Lee, A., Rodriguez-Gonzalez, J., Li, N., Macklin, J.J., Phillips, J.W., Mensh, B.D., Branson, K., Hantman, A.W., 2015. Cortex commands the performance of skilled movement. Elife 4, e10774. https://doi.org/10.7554/eLife.10774

Harvey, C.D., Collman, F., Dombeck, D.A., Tank, D.W., 2009. Intracellular dynamics of hippocampal place cells during virtual navigation. Nature. https://doi.org/10.1038/nature08499

Klaus, A., Martins, G.J., Paixao, V.B., Zhou, P., Paninski, L., Costa, R.M., 2017. The Spatiotemporal Organization of the Striatum Encodes Action Space. Neuron 95, 1171–1180.e7. https://doi.org/10.1016/j.neuron.2017.08.015

Lak, A., Costa, G.M., Romberg, E., Koulakov, A.A., Mainen, Z.F., Kepecs, A., 2014. Orbitofrontal cortex is required for optimal waiting based on decision confidence. Neuron. https://doi.org/10.1016/j.neuron.2014.08.039

Maeda, R.S., Cluff, T., Gribble, P.L., Pruszynski, J.A., 2018. Feedforward and feedback control share an internal model of the arm’s dynamics. J. Neurosci. https://doi.org/10.1523/JNEUROSCI.1709-18.2018

Mathis, A., Mamidanna, P., Cury, K.M., Abe, T., Murthy, V.N., Mathis, M.W., Bethge, M., 2018. DeepLabCut: markerless pose estimation of user-defined body parts with deep learning. Nat. Neurosci. https://doi.org/10.1038/s41593-018-0209-y

Mathis, M.W., Mathis, A., Uchida, N., 2017. Somatosensory Cortex Plays an Essential Role in Forelimb Motor Adaptation in Mice. Neuron 93, 1493–1503.e6. https://doi.org/10.1016/j.neuron.2017.02.049

Osborne, J.E., Dudman, J.T., 2014. RIVETS: A mechanical system for in vivo and in vitro electrophysiology and imaging. PLoS One. https://doi.org/10.1371/journal.pone.0089007

Panigrahi, B., Martin, K.A., Li, Y., Graves, A.R., Vollmer, A., Olson, L., Mensh, B.D., Karpova, A.Y., Dudman, J.T., 2015. Dopamine Is Required for the Neural Representation and Control of Movement Vigor. Cell 162, 1418–1430. https://doi.org/10.1016/j.cell.2015.08.014

Paninski, L., 2003. Spatiotemporal Tuning of Motor Cortical Neurons for Hand Position and Velocity. J. Neurophysiol. 91, 515–532. https://doi.org/10.1152/jn.00587.2002

Robie, A.A., Seagraves, K.M., Egnor, S.E.R., Branson, K., 2017. Machine vision methods for analyzing social interactions. J. Exp. Biol. https://doi.org/10.1242/jeb.142281

Slutzky, M.W., Jordan, L.R., Bauman, M.J., Miller, L.E., 2010. A new rodent behavioral paradigm for studying forelimb movement. J. Neurosci. Methods 192, 228–232. https://doi.org/10.1016/j.jneumeth.2010.07.040

Tai, L.H., Lee, A.M., Benavidez, N., Bonci, A., Wilbrecht, L., 2012. Transient stimulation of distinct subpopulations of striatal neurons mimics changes in action value. Nat. Neurosci. 15, 1281–1289. https://doi.org/10.1038/nn.3188

Thoroughman, K.A., Shadmehr, R., 1999. Electromyographic correlates of learning an internal model of reaching movements. J. Neurosci. 19, 8573–8588.

Yttri, E.A., Dudman, J.T., 2016. Opponent and bidirectional control of movement velocity in the basal ganglia. Nature 533, 402–406. https://doi.org/10.1038/nature17639

Yttri, E.A., Liu, Y., Snyder, L.H., Goldberg, M.E., 2013. Lesions of cortical area LIP affect reach onset only when the reach is accompanied by a saccade, revealing an active eye–hand coordination circuit. PNAS 110, 2371–2376. https://doi.org/10.1073/pnas.1220508110

